# Identification of diverse viruses associated with grasshoppers unveils phylogenetic congruence between hosts and viruses

**DOI:** 10.1101/2021.11.16.468783

**Authors:** Yao Xu, Jingyi Jiang, Xiaoju Lin, Wangpeng Shi, Chuan Cao

## Abstract

Locusts and grasshoppers are one of the most dangerous agricultural pests. Environmentally benign microbial pesticides are increasingly desirable for controlling locust outbreaks in fragile ecosystems. Here we use metagenomic sequencing to profile the rich viral communities in 34 grasshopper species and report 322 viruses, including 202 novel species. Most of the identified viruses are related to other insect viruses and some are targeted by antiviral RNAi pathway, indicating they infect grasshoppers. Some plant/fungi/vertebrate associated viruses are also abundant in our samples. Our analysis of relationships between host and virus phylogenies suggests that the composition of viromes is closely allied with host evolution, and there is significant phylogenetic relatedness between grasshoppers and viruses from *Lispiviridae, Partitiviridae, Orthomyxoviridae, Virgaviridae* and *Flaviviridae*. Overall, this study is a thorough exploration of viruses in grasshoppers and provide an essential evolutionary and ecological context for host-virus interaction in Acridoidea.

**Author Summary:** Locusts are the most destructive migratory pest in the world and continue to cause massive damages that endanger food security and threaten millions of people in 21^st^ century. While chemical pesticides are still heavily relied on, biological pesticides developed from natural pathogens offer a reliable, less harmful alternative for controlling locust outbreaks in fragile ecosystems. Unfortunately, little is known about natural pathogens infecting this pest. In this study, we profile the viral communities in 34 grasshopper species include some major swarming species. While we identified as many as 202 novel viral species associated with grasshoppers, some of them are of potential to be developed as biocontrol agents. Our analysis of relatedness of phylogenies of grasshoppers and associated viruses helps to shed light on the eco-evolutionary interactions between insects and viruses. This work provides a valuable dataset of both academic and applied interest.

## Introduction

Locusts are grasshoppers that can form swarms and migrate long-distance. They are one of the most devastating threats to agriculture throughout human history. Even in the 21^st^ century, locusts still cause massive damages that endangering food security and threatening millions of people [1]. There are currently more than 500 documented species of acridids (Orthoptera: Acridoidea) that can cause damage to pastures and crops, and about 50 are considered major pests [2]. Recent event of desert locusts (*Schistocerca gregaria*) swarms in Arabian Peninsula, East Africa, India and Pakistan since late 2019 were the worst upsurge seen in last seventy years for some countries [3]. Meanwhile, local outbreaks of the Moroccan locust (*Dociostaurus maroccanus*) in Central Asian countries, the Italian locust (*Calliptamus italicus*) in Russia and China, the South American locust (*Schistocerca cancellata*) in Paraguay and Argentina, the African migratory locust (*Locusta migratoria migratorioides*) in southern African countries as well as the Yellow-spined bamboo locust (*Ceracris kiangsu*) in Laos, Vietnam and China also cause major economic, social and environmental impacts [4].

Most grasshopper and locust control still rely on chemical pesticides, and this has raised many issues about human health, environment, non-target organisms and biodiversity [2]. In recent years, there are an increased use of alternative biological control methods. Biopesticides such as *Metarhizium acridum* and *Paranosema locustae* have been used in concerted controlling of grasshoppers and locusts in China and have successfully prevented migratory locusts reaching plague proportions [2, 5]. *M. acridum* has also been used in East Africa against recent desert locust infestation [6] and used operationally against the migratory locust in East Timor and against red locusts in East Africa [2]. Another promising control method is using entomopathogenic viruses, which are environmentally benign, species-specific and can spread horizontally and transmit vertically. However, to date, only a few viruses have been isolated and characterized in grasshoppers: entomopoxviruses in *Melanoplus sanguinipes* and in *Oedaleus senegalensis* [7]-9] and a picornavirus in *Schistocerca americana* [10]. *Melanoplus sanguinipes entomopoxvirus* (MSEV) has been investigated for their potential use as biological control agents against orthopteran insects as they infect many of major grasshopper and locust pest species [11-12]. However, the slow occurring mortality has limited its broad use as microbial insecticides [13]. Thus, there is still a demand in discovering new viral pathogens that could be harnessed to control grasshoppers and locusts.

Apart from discovering novel viral biocontrol agents, understanding the nature of viral infections in grasshoppers may help to better understand the physiology, geographical establishment and evolution of these notorious pests. Recent metatranscriptomic studies of a variety of insect species have revealed that they harbor an enormous diversity of RNA viruses [14-17]. Characterizing viromes with known hosts not only provides a better perspective on the taxonomy and evolution of viruses [18-19], but also sheds light on host-association and host-switching of viruses [20-21]. More and more evidence has shown that the host lineage poses great influence on the composition as well as sequence divergence of virome, and viruses tend to jump between phylogenetically related host species [22-24]. Understanding host-switching/sharing of viruses will be of potential importance for biocontrol decisions in the future.

Orthopteran insects are an underrepresented groups in virome studies with reported viruses belonging to *Flaviviridae, Virgaviridae, Narnaviridae* and *Partitiviridae* in pan-arthropod virome studies [14, 16]. Here, we use a metagenomic approach to characterize the virome associated with 34 species of grasshoppers including many major agricultural pest species, with the emphasis on better characterizing the diversity and abundance of viruses and understanding their eco-evolutionary relationship with hosts.

## Results

### Abundant and divergent viruses identified in grasshoppers

Ten ribosome-depleted total RNAs extracted from six species from three locations were sequenced, generating 8.69 Gb to 15.59 Gb sequence data for each sample. For five of the species, 21.77-94.05% of reads can be mapped to viruses (S1 Table). For *Locusta migratoria*, only 3.29% of the reads mapped to viral-like sequences and majority of reads cannot be mapped at all. Additional selected publicly available transcriptomic data of 28 species of grasshoppers collected across the world were also included in the analysis (S1 Table). These data sets were generated from mashed whole grasshoppers or specific tissues, and total bases obtained varies between 2.3 Gb to 13.8 Gb.

Through a BLASTX search with assembled sequences, a total of 694 candidates viral contigs were identified. Using identifiable RdRp sequences (>200aa), we were able to assign 322 distinctive putative viruses to 44 virus families or unclassified viruses (Fig 1A, S2 Table). The number of viruses identified in each host species varied a lot, with *Chorthippus albonemus* collected from Qinghai containing as many as 47 viruses. Significant fewer viruses were identified in publicly available data sets, possibly because they were lab insect cultures and had smaller sequencing depth (Fig 1B).

**Fig 1.**
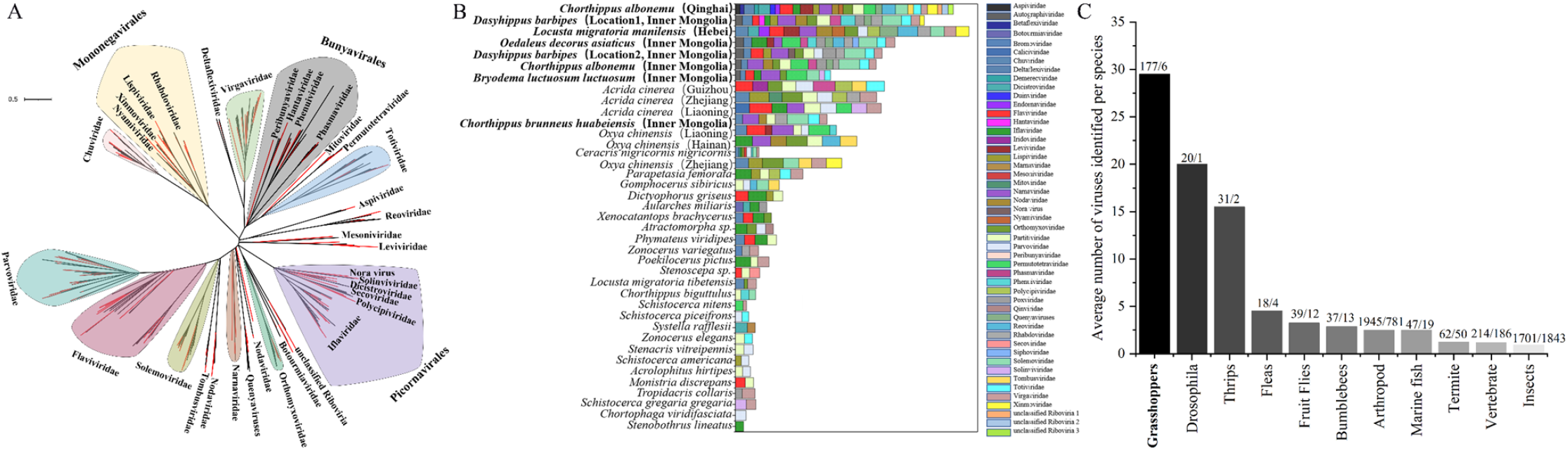
Viruses associated with grasshoppers. (A) ML phylogeny of major RNA viruses. Viruses discovered in this study are in red. (B) Virome composition of 34 grasshopper species. The species marked in bold are metatranscriptional sequenced samples from present study. The different collection locations of the same species are recorded in brackets. (C) Average number of viral species identified in different hosts by metatranscirptomic sequencing studies. The number of viruses and host species are marked above the bars.

In total, 106 positive single-stranded RNA (+ssRNA) viruses, 61 negative single-stranded RNA (-ssRNA) viruses, 48 double-stranded RNA (dsRNA) viruses, 10 double-stranded DNA (dsDNA) virus and 17 single-stranded DNA (ssDNA) viruses were identified in our study (S2 Table). RNA viruses were the dominant type, comprising up to 89% of the identified viruses. Picorna-like viruses including viruses mainly from *Iflaviridae, Dicistroviridae, Solinviviridae* and *Polycipiviridae* were present in 19 grasshopper hosts, accounting for 29% of +ssRNA viruses (Fig 1A, S2 Table). Mononega-like viruses and bunya-like viruses were the most common -ssRNA viruses identified, representing up to 57% of all -ssRNA viruses identified and present in more than 44% grasshopper hosts in this study. Partiti-like viruses found in 47% grasshopper hosts and was the most common dsRNA virus. entomopoxviruses were the only eukaryotic dsDNA virus identified and were found in five grasshopper hosts. 17 parvo-like ssDNA viruses were the only ssDNA virus identified and presented in 12 grasshopper hosts (S2 Table).

The majority of newly identified viruses were highly divergent from previously reported viral sequences: 71% viruses shared less than 50% amino acid (aa) identity with their most closely related RdRp sequence (S2 Table). Based on ICTV species demarcation criteria (https://talk.ictvonline.org/ictv-reports), 202 viruses can be considered as novel species (details in S2 Table). This is a large number of novel viruses identified when compared to similar studies of other organisms (Fig 1C) and has substantially enriched the number of recorded orthopteran viruses [15, 19-20, 25-30]. Novel viruses found in this study are named after their host species, related virus family like, followed by a number (*e*.*g. Chorthippus albonemus* chu-like virus 1). If one virus infects more than one host species, genus or family name of multiple hosts were used (*e*.*g*. Gomphocerinae chu-like virus). Complete or near-complete genome sequences were obtained in 65 novel viral species belonging to 22 families (S1 Data) and tentative genome structures were show in Figure 2. Viruses from the same family tend to share similar genome structure with exceptions of *Virgaviridae, Flaviviridae* and *Totiviridae* (Fig 2). *Virgaviridae* showed a great flexibility in genome size and arrangement. *Flaviviridae* contains both typical segmented genome and a substantially larger unsegmented genome [31]. Toti-like virus in grasshoppers could either encode a capsid protein or a novel proline-alanine rich protein, as described in a previous study [32].

**Fig 2.**
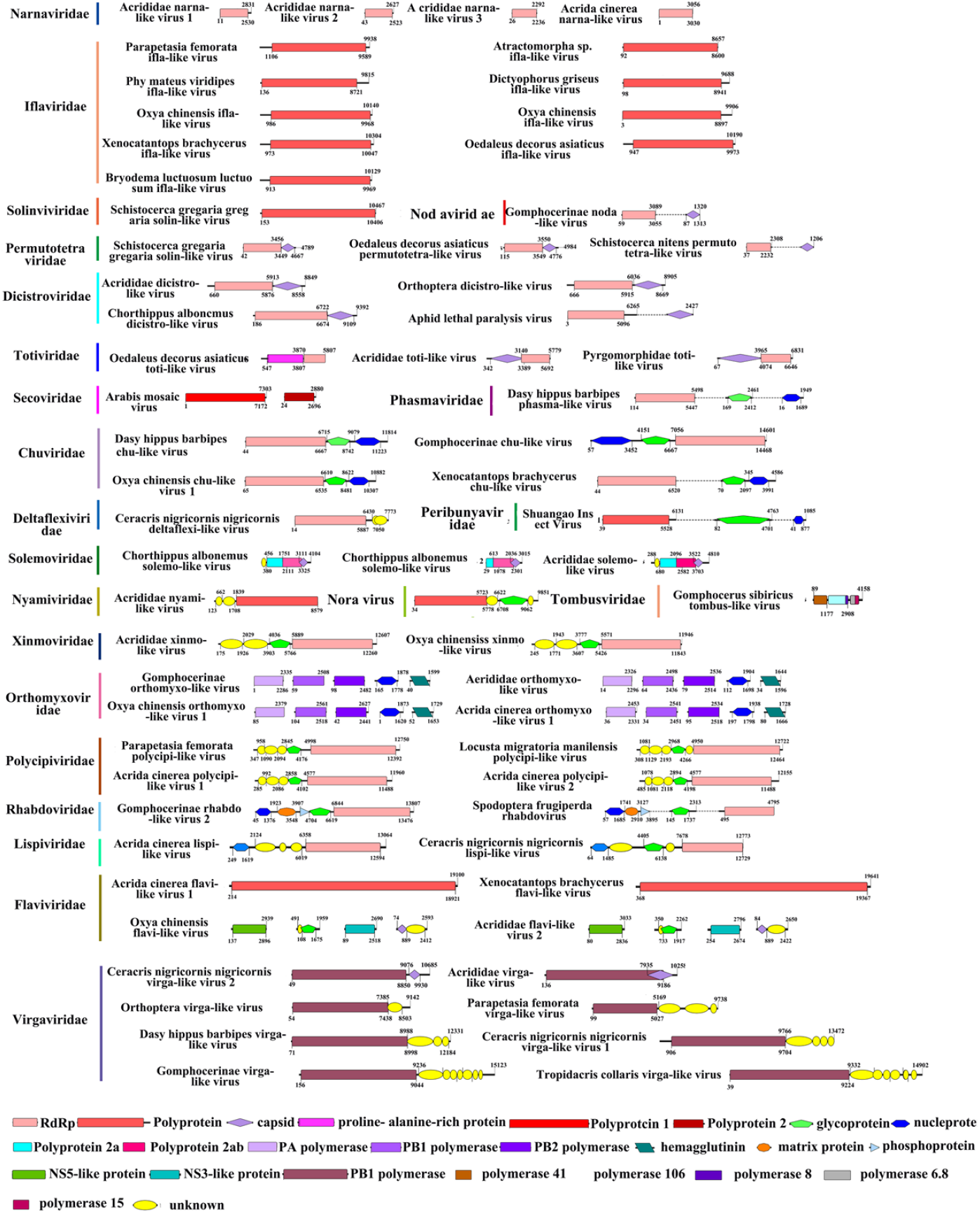
Annotations of complete or near-complete viral genomes. Viral proteins are colored according to their putative functions. For unsegmented viruses, the different functional proteins that lack overlapping sequence regions are connected by dotted lines.

### Phylogeny analysis reveals previously unknown insect-associated virus groups

Contigs of 158 RdRps or conserved domains were grouped, and phylogenetic trees were generated following optimization of alignments. Thirteen trees were generated for 3 viral orders and 29 virus families (Fig 3 and S1-9 Figs). Based on phylogenetic positions in relation to previously described viruses, we sought to make inferences for the hosts of all viruses. 173 identified viruses appear to be insect-associated (S2 Table). Some viruses that have plants or vertebrates as potential primary hosts also exhibit high transcript levels in the analysis such as Acrididae solemo-like virus and Acrididae dicistro-like virus (S3 Table). This indicates that grasshoppers can harbor high copies of plant and vertebrate viruses, which they may acquire through feeding on virus-contaminated food.

**Fig 3.**
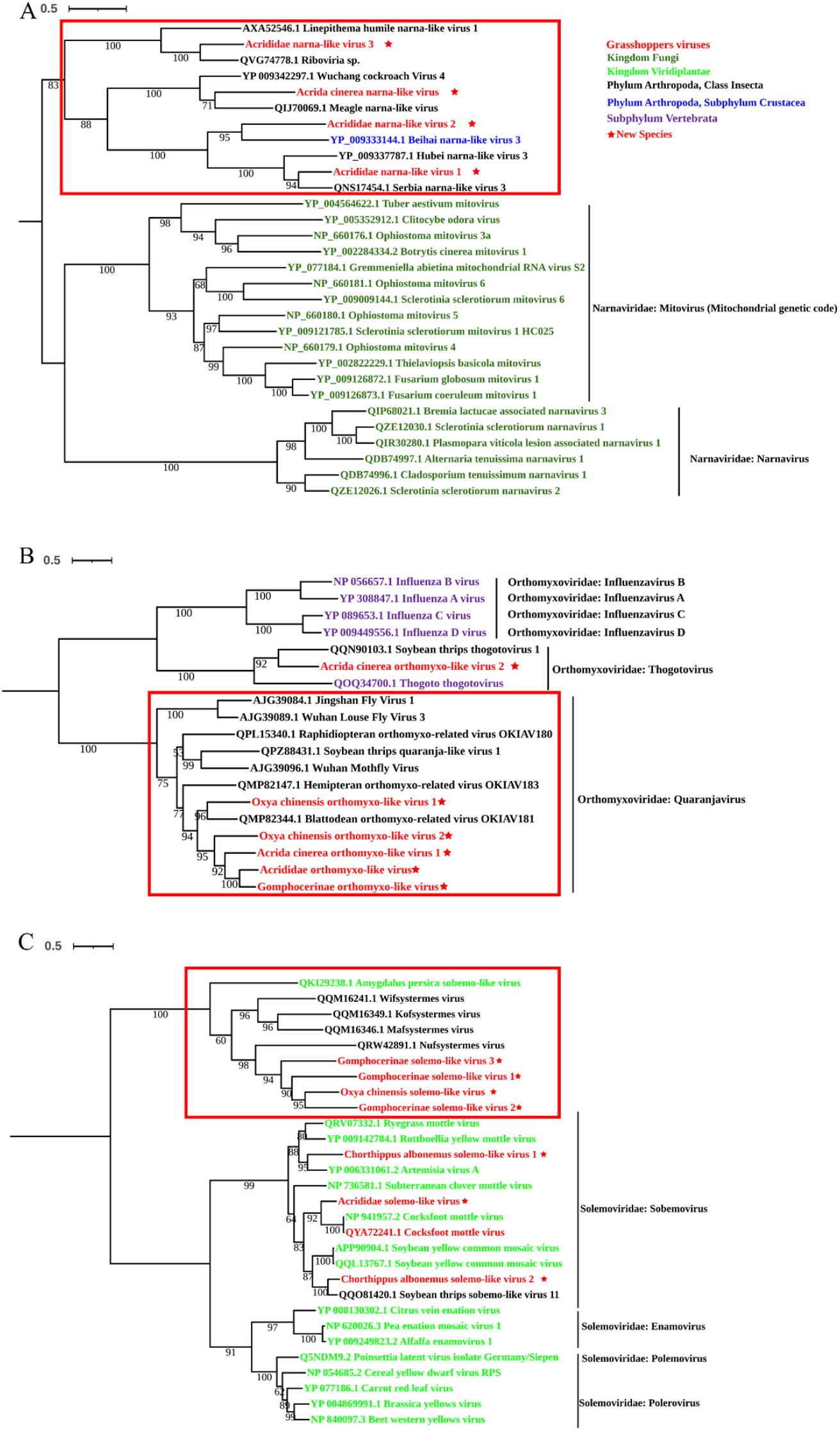
ML phylogenies of *Narnaviridae, Orthomyxoviridae* and *Solemoviridae*. (A) Phylogenetic tree of *Narnaviridae* constructed using RdRp sequences. (B) Phylogenetic tree of *Orthomyxoviridae* constructed using PB1 polymerase. (C) Phylogenetic tree of *Solemoviridae* constructed using replicase sequences. Viruses are colored differently according to their hosts. The viruses described in this study are marked in red and novel viruses have red solid stars at the back of their names.

Interestingly, we found certain viral families that were considered only infecting plant/fungus/vertebrate hosts had formed a separate clade containing viruses discovered in arthropods/insects. For example, four novel narna-like viruses found in this study formed a distinct clade together with viruses discovered in Linepithema ants, cockroaches and other arthropods (Fig 3A); five novel orthomyxo-like viruses grouped together and formed a separate clade with viruses found in flies, thrips and other blattodean and hemipteran insects (Fig 3B); four novel solemo-like viruses grouped together with viruses found in termites (Fig 3C). These insect/arthropod associated virus clades may expand the host range and help to fill evolutionary gaps within these virus families that were previously thought to be plant/fungus/vertebrate specific.

### Phylosymbiosis detected between grasshoppers and their viruses

Although environmental factors are considered to play an essential role, host genetics and evolutionary history may also affect the composition of host virome. If hosts influence a sufficient amount of the composition of virome, then hosts with greater genetic divergence may exhibit more distinguishable viral composition [33-34]. We constructed host phylogeny of ten grasshopper species based on amino acid alignments of the mitochondrial genome and it was consistent with Song et al., (2020) [35]. The viral dendrogram was generated from Bray-Curtis beta diversity of the viral metagenomes of corresponding host species. Host phylogeny showed significant congruence with the branching pattern of the viral dendrogram (*I*_*cong*_=1.46, *p*<0.01) (Fig 4). This result suggests a phylosymbiotic relationship between grasshopper host and viral beta diversity, meaning that evolutionary changes in the host are associated with ecological changes in the virome [33].

**Fig 4.**
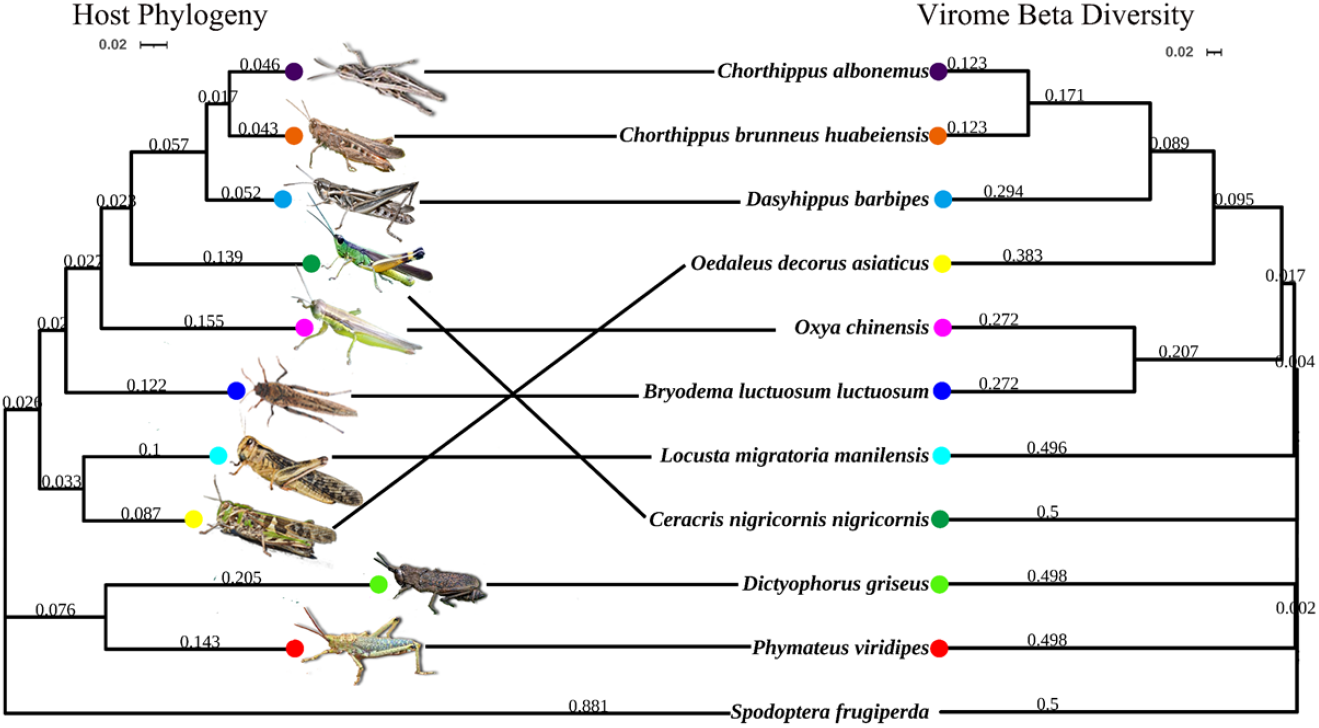
Phylosymbiosis between ten grasshopper species and their viral communities. The host phylogeny is constructed based on the cytochrome oxidase I gene, and the UPGMA hierarchical cluster relationships of the viromes are based on Bray-Curtis beta diversity distances. The topological similarity and significance between the host phylogeny and the virome clustergram was determined by calculating a congruence index described in De Vienne et al., (2007) [36] (*Icong*=1.46, *p*<0.01). Horizontal lines connect species whose position is concordant between host phylogeny and virome clustergram. *Spodoptera frugiperda* and its virome data were used as outgroups for the analysis.

We used a modified Mantel permutation test [37-38] to test if the phylogenetic tree of a virus family is related to the phylogenetic tree of their hosts. The host phylogeny was constructed for 32 species that have near-complete mitochondrial genome. The topology of the phylogeny of virus families were compared with that of their hosts using their pairwise patristic distances. Results showed that the patristic distances of families *Lispiviridae* (*r*=0.88, *P*<0.05), *Partitiviridae* (*r*=0.62, *P*<0.005), *Orthomyxoviridae* (*r*=0.44, *P*<0.05), *Virgaviridae* (*r*=0.38, *P*<0.05) and *Flaviviridae* (*r*=0.31, *P<*0.05) were significantly correlated with their host patristic distance. This congruence of virus and host phylogenies indicates that these viruses may coevolve with their primary grasshopper hosts.

### Natural prevalence and cross-species transmission of grasshopper viruses

To examine if some viruses can infect diverse hosts, we applied PCR tests on multiple natural populations of grasshoppers collected in year 2018 and 2021. Five species of grasshoppers were collected from Qinghai in 2018 and only three viruses out of 53 tested could be detected in some Qinghai populations. The natural infection rates of these three viruses were provided in Table 1.

**Table 1.**
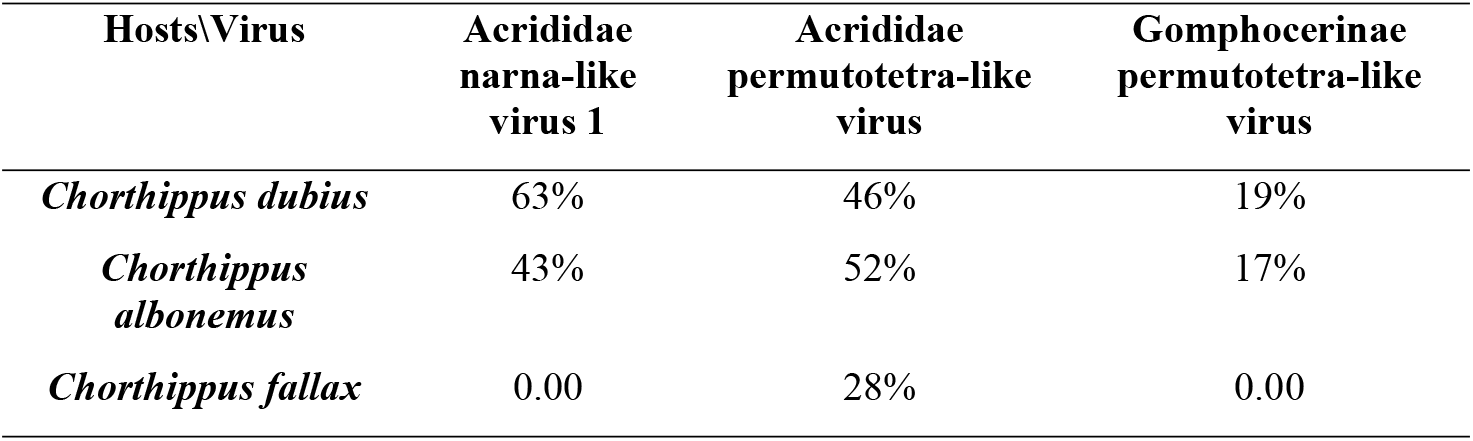
The average natural infection rates of viruses in the wild.

| Hosts\Virus | Acrididae<br>narna-like<br>virus 1 | Acrididae<br>permutotetra-like<br>virus | Gomphocerinae<br>permutotetra-like<br>virus |
| --- | --- | --- | --- |
| *Chorthippus dubius* | 63% | 46% | 19% |
| *Chorthippus albonemus* | 43% | 52% | 17% |
| *Chorthippus fallax* | 0.00 | 28% | 0.00 |

We also collected *Dasyhippus barbipes* and *Bryodema luctuosum luctuosum* in 2019 and 2021 separately. Through PCR tests, 67% of the viruses identified in 2019 were found in *B. luctuosum luctuosum* collected in 2021, while only 22% of viruses identified in 2019 were detected in *D. barbipes* grasshoppers in 2021 (S4 Table).

Co-occupying the same ecological niches may facilitate cross-host transmission of certain viruses. Viruses from families *Naranaviridae* and *Permutotetraviridae* tend to infect multiple host species (S2 Table). For example, Acrididae narna-like virus 1 was present in seven host species, Acrididae narna-like virus 3 was present in four host species, Acrididae permutotetra-like virus was present in five host species. This result indicates that viruses from these families may be transmitted more often across different host species.

### Antiviral RNAi against various viruses in *Locusta migratoria*

Different from intensively studied RNAi response in *Drosophila melanogaster*, previous study did not find typical siRNA 21nt peak nor piRNA pattern in the distribution of virus-derived siRNAs (vsiRNAs) in *L. migratoria* [39]. Using Lewis’ dataset (SRA: SRS2228471, SRS2228473) [39], we identified contigs of seven DNA viruses and one +ssRNA virus, including five insect-specific viruses: a granulovirus, an entomopoxvirus, an iridovirus and two nucleopolyhedraviruses. After filtering host genome sequence, oxidation-treated sRNAs were mapped to viral contigs and interestingly, seven of them (except for a virga-like virus) showed a 22nt peak in their distributions (Fig 5A). This indicates that *L. migratoria*’s antiviral RNAi pathway may generate 22nt biased sRNA for some DNA and RNA viruses.

**Fig 5.**
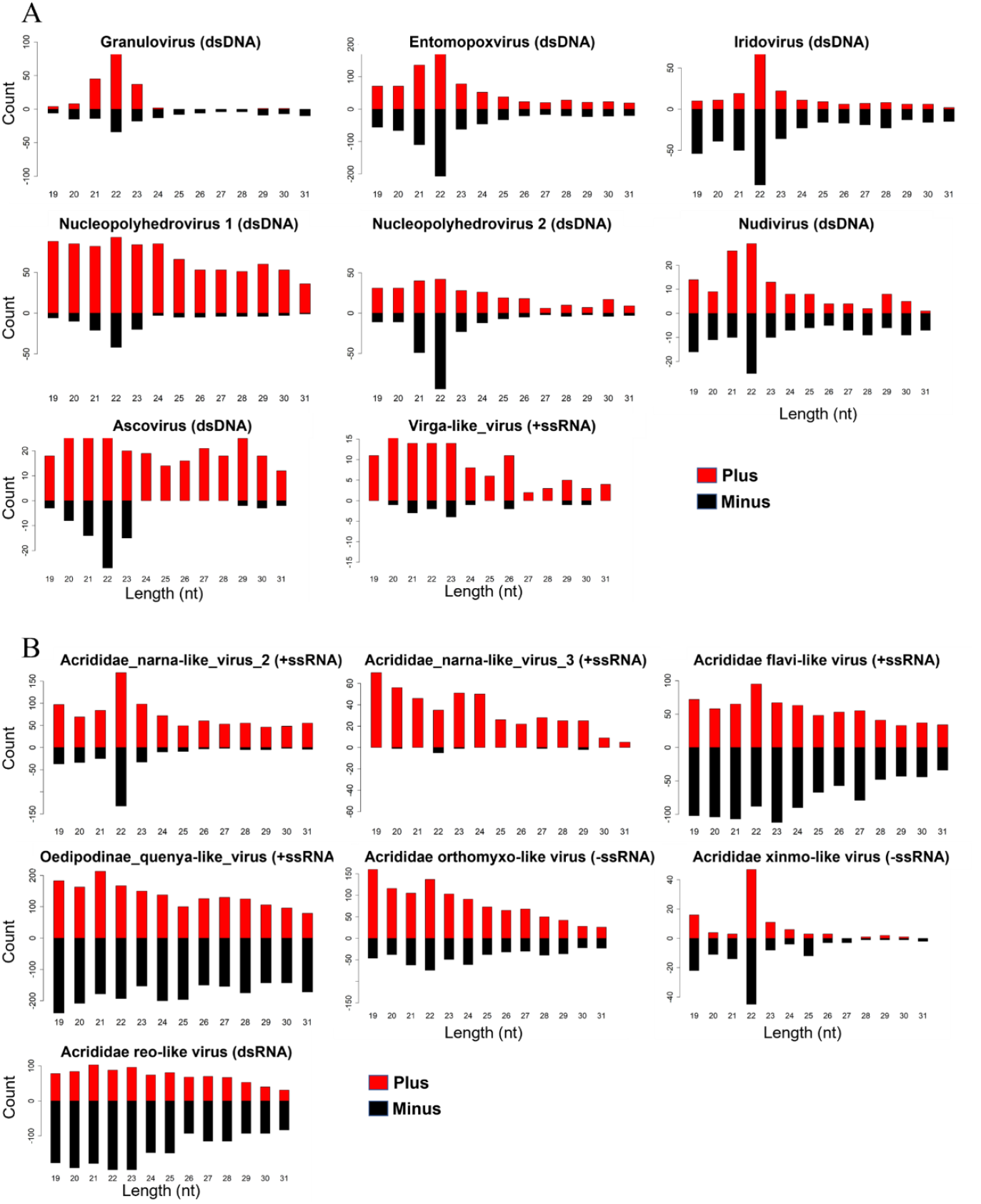
Small RNA size distribution. (A) Small RNA distribution of eight viruses identified using data from Lewis et al., (2018) [39]. (B) Small RNA distribution of seven viruses identified in this study.

To further explore siRNA-based antiviral immunity in grasshoppers, we carried out small RNA sequencing on the same *L. migratoria* samples which we used for virus RNAseq. Among 25 viruses that were found in *L. migratoria*, sRNAs were successfully mapped to contigs of seven viruses after filtering out host genome sequences. sRNAs mapped to Acrididae narna-like virus 2, 3 and Acrididae xinmo-like virus also showed an obvious enrichment in 22nt (Fig 5B). Other five viruses including three +ssRNA viruses, one -ssRNA virus and one dsRNA virus, did not show obvious 21nt nor 22nt peak (Fig 5B). In either dataset, we did not find virus-derived piRNAs that bearing the signature of ping-pong amplification. Notably, for many viruses such as Oedipodinae noda-like virus, Acrididae solemo-like virus, Drosophila A virus that have high abundance in the host (S3 Table), no sRNA was found mapped to them. These results suggest that antiviral RNAi pathways is actively involved in response to viruses and the distribution of sRNA may vary for different viruses.

## Discussion

In this study, we present the first survey of virome of a notorious insect pest, grasshoppers, and demonstrate that they harbor a diverse range of viruses. Overall, 215 RNA viruses and 27 DNA viruses were identified. These viruses are quite divergent from previous known species, and 202 of them can be considered as novel species. Although the potential of these viruses be used as biological control agents is currently unclear, there are some good candidates. For example, entomopoxvirus and densovirus have been registered as biocontrol agents [40] and we identify three novel entomopoxviruses and 17 novel densoviruses infecting five and twelve host species separately. Genes of another identified virus, iridovirus, are explored as biocontrol compounds for their toxicity effect on insect hosts [41]. Reovirus which was recorded causing epizootics in natural populations of insects was also found in six grasshopper species in this study [42]. Note that sequences of baculovirus and granulovirus which are broadly used biopesticides were also found when we analysed Lewis’ data [39], but they are not present in our data. Further isolation and pathogenicity assays are required to evaluate the potential use of these viruses as control agents.

Among all identified viruses, 71% of them are believed to be insect-associated based on virus phylogenies. Another 39 plant/fungus-associated viruses and 8 vertebrate-associated viruses were identified. Grasshoppers may ingest these viruses from contaminated food. It is not clear if these viruses can infect and replicate in grasshoppers with one exception: we are able to find Acrididae solemo-like virus (a plant virus) in *L. migratoria* infected with crude virus extract but not in the control (unpublished data). However, high abundance of some of these plant and vertebrate viruses in grasshoppers provide a possibility that viruses may transmit from insects back to plant and animal hosts by contacting or feeding [43-45]. Thus, grasshoppers are potential vectors of plant and vertebrate viruses and may facilitate transmission of these viruses.

Whether host genetics play an essential role in shaping the virome composition and in the evolution of viruses remains an intriguing question. Of the ten grasshopper species that were analysed, significant topology congruence between host phylogeny tree and virus Bray-Curtis beta dendrogram were found. This suggests that in natural environment, phylosymbiotic relationship may exist between these grasshoppers and their virome. In addition, we find significant relatedness between host phylogeny and phylogenies of Lispi-like, Partiti-like, Orthomyxo-like and Flavi-like viruses. These virus families are either recently proposed monogenetic families or contain newly discovered insect-specific clades [28, 31]. Many other studies have found phylosymbiosis between hosts and microbial communities (mostly bacteria and fungi) in a diverse range of systems under controlled regimes [33, 46] and under natural environment [47-48]. In certain plant [49] and animal [50] systems, significant phylosybmbiosis were also found between hosts and virus communities. Our results highlight that host phylogeny is significantly associated with its virome composition and virus evolution. However, we need to be cautious about these conclusions for 1) there is considerable topological uncertainty in virus phylogenies and 2) functional studies of both hosts and viruses are required. Nevertheless, with better characterizing of viruses associated with wider insect hosts, we will have a better chance of understanding the eco-evolutionary relationships between hosts and viruses.

By co-analysis of metagenomics and sRNA data, we are able to show that the antiviral RNAi may play an essential role in defense against viruses in *L. migratoria*. The virus-derived interfering RNA (viRNA) profile of *L. migratoria* shows a 22nt peak for both DNA and RNA viruses. Similar viRNAs distributions were observed in other insects, such as in whiteflies [24], thrips [29] and bumblebees [27]. It is possible that the 21nt peak we seen in brachycera species such as flies and mosquitoes is unusual for insects [51-52].

Like all other metagenomics studies, our work has several limitations [53]. For instance, our virus identification purely based on sequence homology search. With high divergence, only the most conserved sequences are recognizable at the protein level. Indeed, for many novel viruses found in this study, especially those with segmented genomes, we had difficulty in identifying other proteins besides the RdRp. Moreover, we had difficulty in determining if these identified viruses are grasshopper-infecting. Additional sRNA sequencing would be useful in solving this issue in the future. Nevertheless, this study is a significant addition to our understanding of the abundance and diversity of insect-associated viruses and their molecular, evolutionary interactions with insect hosts, providing a rich resource for developing biological control agents for controlling grasshopper and locust pests.

## Materials and methods

### Sample collection and virus isolation

Grasshoppers were collected by sweep-netting grassland from ten locations in Inner Mongolia and Qinghai, China between 2018 and 2021, with an average of 618 individuals per species (S1 Table). *Locusta migratoria* were purchased from a grasshopper breeding center, Hebei, China. Species were identified using morphological characteristics and mitochondrial cytochrome c oxidase subunit I gene (COI) sequences.

Crude virus purification was performed for *C. albonemus, Chorthippus brunneus huabeiensis, D. barbipes, B. luctuosum luctuosum, Oedaleus decorus asiaticus* collected in Inner Mongolia, *C. albonemus* collected in Qinghai, and *L. migratoria*. Briefly, pools of grasshoppers of the same species were homogenized in Ringer’s solution and debris removed by low-speed centrifugation at 500 × g for 10 min. The supernatant was layered on top of a discontinuous sucrose gradient (30%, 40%, 50%, 60% w/v) and centrifuged at 64,000 x g for 3 hours in A27-8×50 mL rotor (Beckman Coulter). The visible virus bands were collected, mixed, and centrifuged for 2 h at 64,000 g in A27-8×50 mL rotors (Beckman Coulter) to sediment virus particles. Viral particles were then suspended in 500 uL of DNase/RNase-Free water (Solarbio).

### RNA sequencing and reads assembly

Total viral RNA was extracted using TRIzol LS reagent (Invitrogen, Carlsbad, CA) according to the manufacturer’s instructions. Viral RNA quality was examined using NanoDrop 2000 (ThermoFisher Scientific, Waltham, MA) and Agilent 2100 bioanalyzer (ThermoFisher Scientific, CA, USA). Residual Ribosomal RNA (rRNA) was depleted using Ribo-Zero™ kits (Epicentre, Madison, WI) before library construction. Libraries were constructed using a TruSeq total RNA library preparation kit (Illumina) and paired-end (250-300bp) sequencing was performed on the Hiseq-PE150 platform (Illumina, Sandiego, CA). Additionally, RNA-Seq data of 28 grasshoppers collected worldwide were retrieved from the NCBI database (S1 Table). Sequencing reads were quality trimmed using trimmomatic and *de novo* assembled using the Trinity version 2.8.6 with default parameter settings and the minimum contig length was set at 200nt [54].

### Discovery of viral sequences

The assembled contigs were compared to reference viral protein database (taxid: 10239) downloaded from NCBI using Diamond BLASTx version 2.0.11 [55], with e-value cut-offs of 1 × 10^−5^. To eliminate false positives, the putative viral contigs were then compared to the entire non-redundant protein database (nr) of NCBI using Diamond BLASTx version 2.0.11 [55]. Contigs with credible, significant BLAST hits (e-value < 1 × 10^−5^) to only viral proteins were kept for further analysis. To detect highly divergent viruses, open reading frames were predicted using the open-source NCBI ORF finder (https://www.ncbi.nlm.nih.gov/orffinder/). Predicted amino acid sequence lengths less than 200aa were removed from following analysis. To reduce redundancy, amino acid sequences were grouped based on sequence identity using the CD-HIT package version 4.6.5 [56]. Predicted ORFs without any BLASTx hits were searched for homologous proteins in the protein families (PFAMs) database (http://ftp.ebi.ac.uk/pub/databases/Pfam/releases/Pfam33.1/Pfam-A.hmm.gz) and the RNA-dependent RNA polymerase (RdRp) database of RNA viruses by the use of HMMER version 3.3 [15, 57].

The structure of near complete viral genomes was annotated after comparing them against the entire non-redundant protein database and the genome of the closest virus using BLASTp. To estimate virus abundance in each library, Salmon version 1.4.0 [58] was used to calculate the number of transcripts per million (TPM) of each contig, which was normalized by sequencing depth (total number of reads) and sequence length.

### Phylogenetic analysis

The RdRp or polyproteins of viruses discovered in this study were aligned with sequences of known viruses from same families using MAFFT version 7.158 with the E-INS-i algorithm [59]. For a phylogeny tree of major RNA viruses, poorly aligned regions were removed using trimAl version 1.4.1 with a maximum gap threshold of 0.8 and minimum similarity threshold of 0.001 [60]. Phylogenetically informative sites were selected using Gblocks version 0.91b [61]. Maximum likelihood (ML) phylogenetic trees were constructed using IQ-TREE version 1.6.12 with 1,000 bootstraps [62], and the best amino-acid substitution model were determined by Raxml version 2.0 [63]. COI sequences or near-complete mitochondrial genomes of grasshoppers were aligned using MAFFT version 7.158 with the E-INS-i algorithm [59]. Maximum likelihood trees of the grasshoppers were constructed using the same method described above.

### Comparing phylogenies of grasshoppers and viruses

A Modified Mantel permutation test [37-38] was used to test if the phylogenetic tree of each virus family was related to the phylogenetic tree of a host. The host phylogeny was constructed for 32 species that have complete or near-complete mitochondria genome. The topology of the phylogeny of virus families were compared with that of their hosts using their pairwise patristic distances calculated by ParaFit [64] using the R package ape. P-values were calculated from 10,000 iterations of randomized host-virus associations. R version 4.1.1 was used [65].

Virome similarities between hosts were measured by Bray-Curtis distances and employing unweighted pair-group method with arithmetic means (UPGMA) [34]. The R packages Vegan version 2.5-6 [66] was used to calculate Bray-Curtis distances using the virus abundance in each library (TPM) and constructed UPGMA clustergrams between host viromes. The topological similarity and significance between the host phylogeny and the virome clustergram was determined by calculating a congruence index described in De Vienne et al., (2007) [36]. *Spodoptera frugiperda* and its virome data were used as outgroups for the analysis [67].

### Testing prevalence of viruses in natural grasshopper populations

To survey the prevalence of viruses discovered in this study, we examined the presence of 53 virus species in 5 wild grasshoppers’ populations sampled across 9 locations in Qinghai Province, China, with an average of 85 individuals per population (S1 Table). Total virus RNA was prepared from these grasshoppers as described above. Reverse transcription was carried out using PrimeScript™RT reagent Kit (Takara) using random hexamers and PCR was performed with PrimeSTAR Max DNA Polymerase (Takara). Primers were designed based on the assembled viral contigs. We tried to design degenerate primers based on conserved positions for similar viruses and specific primers for poorly conserved virus sequences (S5 Table).

We also checked the prevalence of the identified viruses from *D. barbipes* and *B. luctuosum luctuosum* populations in different years. In summer 2021, we collected grasshoppers in same locations of 2019 and performed PCR for 42 virus species. Specific primers were designed for each of identified viruses from *D. barbipes* and *B. luctuosum luctuosum* populations (S4 Table).

### Small-RNA Sequencing and data analysis

*Locust migratoria* were collected and used for small RNA sequencing. Total RNA was extracted and small RNA sequencing was performed using the high-throughput Illumina nova6000 sequencing technology. Two samples were sequenced 1.194 Gb and 1.292 Gb sRNA data were generated for each sample. The same RNA samples were also used for metatranscriptomic sequencing and virus contigs were obtained used methods above. The sRNA sequences obstained were put through quality control then mapped to *L. migratoria* genome [68] to remove host sequences. Filtered reads were then mapped to various viral contigs using bowtie2 version 2.4.4. The distribution of sRNAs was analyzed using R package viRome [69].

## Acknowledgements

We thank Wang Huiping, Nie Weilei, Ume Hani, Yang Kejia, Tian Jingjing, Feng Mingyue for their assistance in collecting samples.

## Supporting information

**S1 Fig. Maximum likelihood phylogeney of the *Chuviridae, Lispiviridae, Rhabdoviridae, Xinmoviridae*, and *Nyamiviridae***. Phylogenetic tree was constructed using RdRp or polymerase sequences. Viruses were colored differently according to their hosts. Within each phylogeny, viruses described in this study are marked in red and novel viruses have red solid pentagrams at the back of their names.

**S2 Fig. Maximum likelihood phylogeney of the *Caliciviridae, Dicistroviridae, Iflaviridae, Polycipiviridae, Secoviridae*, and *Solinviviridae***. Phylogenetic tree was constructed using RdRp or polymerase sequences.

**S3 Fig. Maximum likelihood phylogeny of the *Hantaviridae, Peribunyaviridae, Phasmaviridae*, and *Phenuiviridae***. Phylogenetic tree was constructed using RdRp or polymerase sequences.

**S4 Fig. Maximum likelihood phylogeny of the *Virgaviridae***. Phylogenetic tree was constructed using RdRp or polymerase sequences.

**S5 Fig. Maximum likelihood phylogeny of the *Partitiviridae***. Phylogenetic tree was constructed using RdRp or polymerase sequences.

**S6 Fig. Maximum likelihood phylogeny of the *Flaviviridae***. Phylogenetic tree was constructed using RdRp or polymerase sequences.

**S7 Fig. Maximum likelihood phylogeny of the *Totiviridae***. Phylogenetic tree was constructed using RdRp or polymerase sequences.

**S8 Fig. Maximum likelihood phylogeny of the *Parvoviridae***. Phylogenetic tree of *Parvoviridae* constructed using capsid protein.

**S9 Fig. Maximum likelihood phylogeny of the *Nodaviridae, Permutotetraviridae, Quenyaviruses, Reoviridae, Tombusviridae*, and *Totiviridae***. Phylogenetic tree was constructed using RdRp or polymerase sequences.

**S1 Table. Sample collection information and data availability**. Excel table providing host species, NCBI project accessions, Collection location and time, Sequencing strategy and depth.

**S2 Table. Identified viruses from this study**.

**S3 Table. Virome composition of 34 grasshopper species and viruses’ abundance**.

**S4 Table. The prevalence of the viruses from *Dasyhippus barbipes* and *Bryodema luctuosum luctuosum* populations in different years**. Excel table providing primer sequences of 42 viruses used to detect the persistence of the viruses.

**S5 Table. Primer sequences of 53 viruses used to investigate natural prevalence. S1 Data. Complete or near-complete genome sequences of 65 novel viral species**.

